# The oxysterol synthesizing enzyme CH25H contributes to the development of intestinal fibrosis

**DOI:** 10.1101/424820

**Authors:** T. Raselli, A. Wyss, M.N. Gonzalez Alvarado, B. Weder, C. Mamie, M.R. Spalinger, W. T. van Haaften, G. Dijkstra, A.W. Sailer, P.H. Imenez Silva, C.A. Wagner, V. Tosevski, Sebastian Leibl, M. Scharl, G. Rogler, M. Hausmann, B. Misselwitz

## Abstract

Intestinal fibrosis and stenosis are common complications of Crohn’s disease (CD), frequently requiring surgery. Anti-inflammatory strategies can only partially prevent fibrosis; hence, anti-fibrotic therapies remain an unmet clinical need. Oxysterols are oxidized cholesterol derivatives, with important roles in various biological processes. The enzyme cholesterol 25-hydroxylase (CH25H) converts cholesterol to 25-hydroxycholesterol (25-HC), which modulates immune responses and oxidative stress. In human intestinal samples from CD patients we found a strong correlation of *CH25H* mRNA expression with the expression of fibrosis markers. We demonstrate reduced intestinal fibrosis in mice deficient for the CH25H enzyme using the sodium dextran sulfate (DSS)-induced chronic colitis model. Additionally, using a heterotopic transplantation model of intestinal fibrosis, we demonstrate reduced collagen deposition and lower concentrations of hydroxyproline in CH25H knockouts. In the heterotopic transplant model, CH25H was expressed in fibroblasts. Taken together, our findings indicate an involvement of oxysterol synthesis in the pathogenesis of intestinal fibrosis.

## Introduction

Crohn’s disease (CD) is a major form of inflammatory bowel disease (IBD), characterized by chronic discontinuous inflammatory lesions. Inflammation in CD is typically transmural and can affect the whole gastrointestinal tract with a preference for the small intestine. Common complications in CD patients include perforations of the gut wall (fistulae and abscesses) as well as intestinal fibrosis and strictures with narrowing of the intestinal lumen. More than 60% of CD patients have to undergo surgery within 20 years following the initial diagnosis[1] and recurrent disease requires more surgical procedures in at least 50% of the patients after the first operation[2, 3]. The second major form of IBD, ulcerative colitis (UC), characterized by continuous inflammatory lesions of the colon, has once been considered a non-fibrotic disease, but recent evidence indicates some degree of submucosal fibrosis in up to 100% of UC colectomy specimens[4, 5] and the degree of fibrosis seems to be proportional to the degree of chronic but not active inflammation[6].

Currently, no drugs have been approved for treatment or prevention of intestinal fibrosis[7, 8]. Anti-inflammatory medications including anti-tumour necrosis factor (TNF) antibodies or immunosuppressants, are only partially effective in preventing fibrosis[9] and new preventive and therapeutic strategies are therefore urgently needed.

On a molecular level, fibrosis is characterized by excessive accumulation of extracellular matrix (ECM) components including collagen and laminin, replacing the original tissue and leading to stiffening and loss of normal function[10, 11]. Transforming growth factor-β (TGF)-β is a key driver of fibrosis, promoting differentiation of fibroblasts to myofibroblast, indicated by expression of α-smooth muscle actin (SMA)[12, 13]. Myofibroblasts are the main effector cells for fibrosis and mainly responsible for ECM deposition[14-16]. On the other hand, myofibroblasts also synthesize matrix metalloproteinases (MMPs) as ECM degrading enzymes and their inhibitors (tissue inhibitor of MMPs, TIMP). Myofibroblasts can derive from the local fibroblast pool; however, epithelial, endothelial, hematopoietic cells, or pericytes can also differentiate into myofibroblasts[16]. Nevertheless, the chain of events leading to intestinal fibrosis is insufficiently understood.

Studying the pathophysiology of intestinal fibrosis has been limited by the lack of a *bona fide* animal model. Chronic dextran sodium sulfate (DSS) colitis is frequently used as a fibrosis model [17, 18], even though key aspects of CD associated intestinal fibrosis, such as occlusion of the intestinal lumen are not observed in this model. Recently, we established and characterized a murine heterotopic transplant model, where small intestinal sections are transplanted into the neck fold of recipient mice[19, 20]. In the transplanted sections, the lumen progressively occludes, accompanied by expression of TGF-β and α-SMA, as well as collagen deposition in the extracellular matrix. In this model, we previously demonstrated that pirfenidone, an anti-inflammatory and anti-fibrotic drug approved for the treatment of idiopathic pulmonary fibrosis, was able to reduce fibrosis[20].

Oxysterols are increasingly recognized as immune-modulatory molecules. 25-hydroxycholesterol (25-HC) is part of the rapid innate immune response and an efficient defence molecule. 25-HC can induce macrophage activation[21-23], T cell differentiation[24], production of IL-8[25-27] as well as IL-6[23] and was shown to have strong antiviral activity against many enveloped viruses[28-30]. Furthermore, Dang and colleagues recently demonstrated a critical role of 25-HC in inhibiting activation of the DNA sensor protein AIM2, preventing spurious AIM2 inflammasome activation[31]. Cholesterol 25-hydroxylase (CH25H) is the key enzyme mediating hydroxylation of cholesterol to 25-HC[32]. 25-HC is rapidly produced *in vivo* upon immune stimulation by toll like receptor (TLR) agonists including lipopolysaccharide (LPS) and poly(I:C)[28, 30, 33, 34]. 25-HC production was shown to be increased in the airways of patients with chronic obstructive pulmonary disease (COPD) and correlated with the degree of neutrophilic infiltration[35]. 25-HC can be further hydroxylated to di-hydroxy cholesterols (e.g. 7α, 25-HC) which have been shown to act as chemoattractants for cells of the adaptive and innate immune system[36, 37]. Recently, CH25H expression was shown to be upregulated in primary lung fibroblasts in response to activated eosinophils, suggesting CH25H activation in chronic lung diseases including COPD[38]. Pro-fibrotic effects of the CH25H product 25-HC have been demonstrated *in vitro*: In a tissue culture model using human fetal lung fibroblasts (HLF), 25-HC induced nuclear factor-κB (NF-κB) activation with subsequent release of TGF-β, leading to myofibroblast formation, MMP-2 and 9 release, SMA expression and collagen production[39]. However, the role of 25-HC in intestinal inflammation and fibrosis has not been addressed. In this study, we aimed to investigate the role of the enzyme CH25H in the development of intestinal fibrosis.

## Results

### *CH25H* mRNA expression is a marker of fibrosis in intestinal samples of CD patients

To test for a role of the oxysterol synthesiting enzyme CH25H in CD associated fibrosis, *CH25H* mRNA expression was measured in human intestinal surgical samples. We investigated terminal ileum samples from CD patients undergoing ileocecal resection due to stenosis. Samples macroscopically affected by fibrosis were compared to the proximal ileal resection margin which showed no macroscopic signs of fibrosis or inflammation. Healthy tissue from cancer-free resection margins of colon adenocarcinoma patients undergoing right-sided hemicolectomy was used as additional control (Table 1). Representative Sirius red staining pictures illustrate increased collagen deposition in fibrotic areas of CD patients (Figure 1A). We observed a gradual increase of *CH25H* mRNA expression from control tissue to non-fibrotic CD tissue and to fibrotic areas from the same patients (p<0.05, Figure 1B). Thereby, mRNA levels of *CH25H* strongly correlated with the expression levels of fibrosis markers including *COL-1* and *-3*, *SMA* and *TGF-β* (Figure 1C-F), confirming the association of CH25H expression with intestinal fibrosis in the human intestine.

**Table 1:**
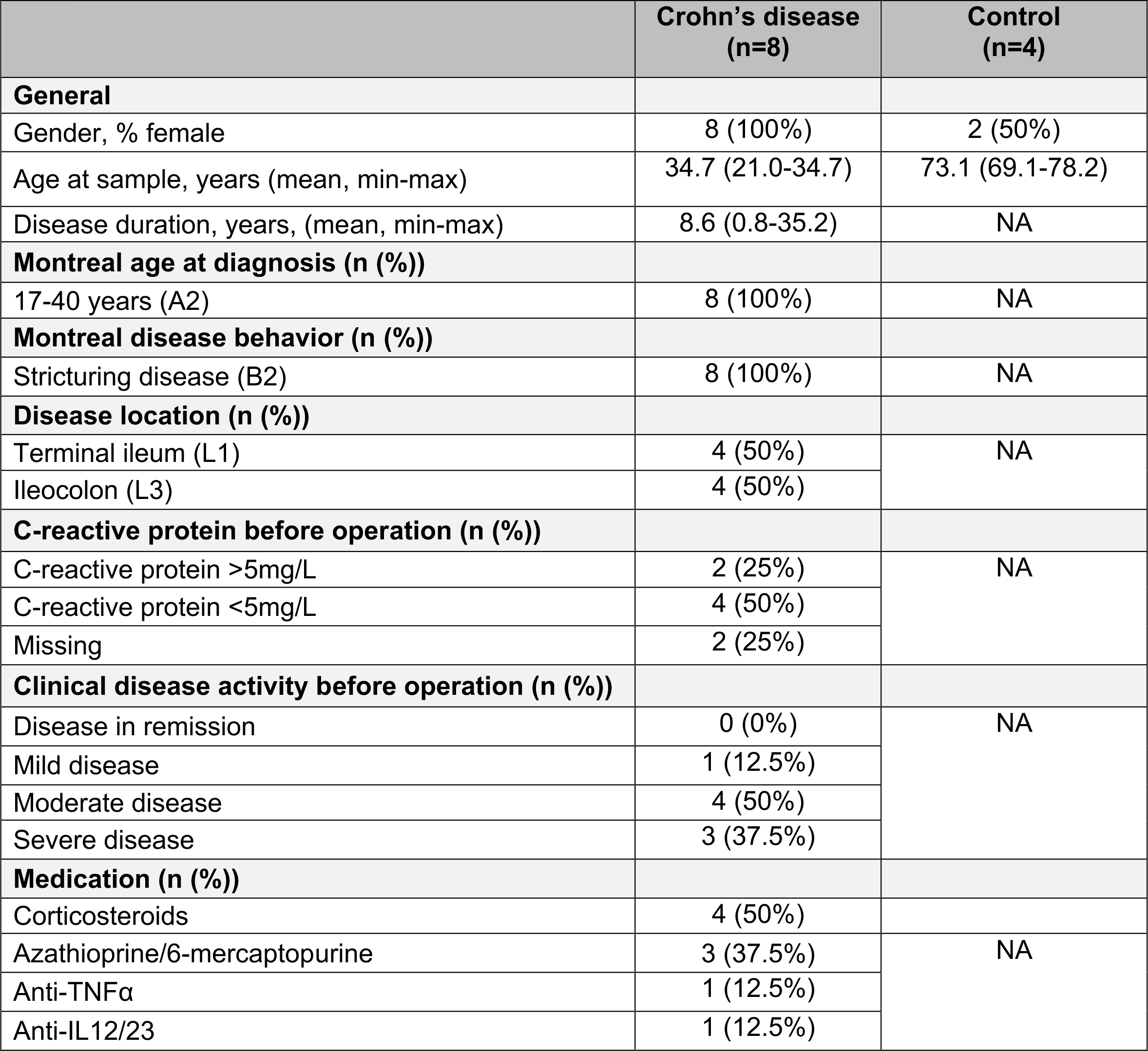
Characteristics of patients with CD and controls. NA: Non-applicable.

**Figure 1:**
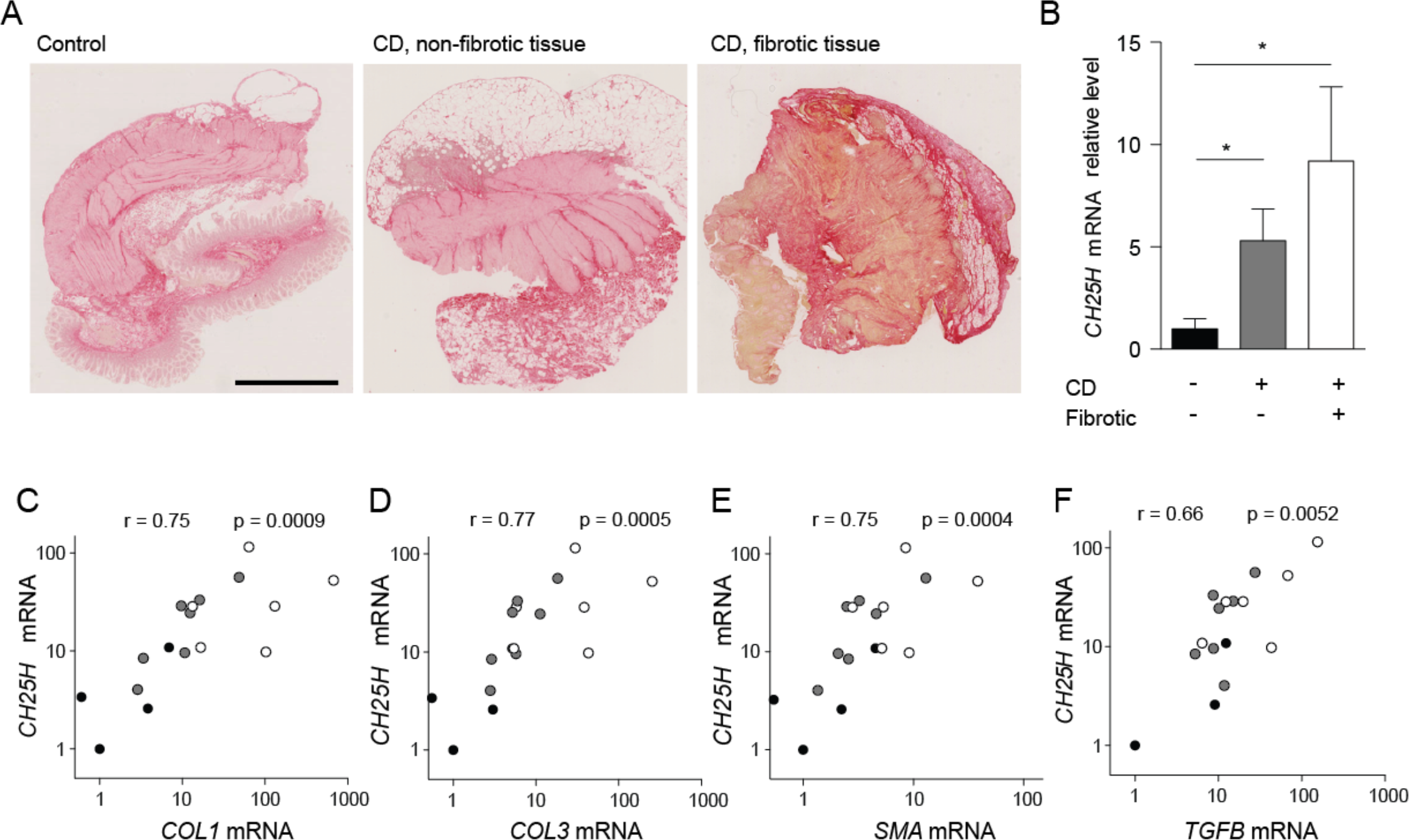
Upregulation of *CH25H* mRNA expression in human fibrotic tissue of patients with CD. (A) Representative images of Sirius-red stained human ileum samples from healthy controls (left panel) and CD patients in a non-fibrotic (middle panel) and in a fibrotic region (right panel). Scale bar: 2.5 mm. **(B)** Samples were analysed for *CH25H* mRNA expression and normalized to *GAPDH*. *CH25H* mRNA level was correlated with mRNA levels of **(C)** *COL1*, **(D)** *COL3* **(E)** *SMA* and **(F)** *TGFB*. White: CD fibrotic, (n=6), grey: CD non fibrotic (n=7), black: healthy control (n=4). Statistical analysis: B: Mann-Whitney U test; * = p < 0.05. CD, Crohn’s disease. C-E: Correlation analysis: Spearman R (non-parametric correlation).

### Reduced intestinal fibrosis in mice with deficient 25-hydroxycholesterol synthesis

To further investigate the role of CH25H in intestinal fibrosis, we investigated whether absence of CH25H would reduce fibrosis in dextran sodium sulfate (DSS)-induced chronic colitis, a well-established model of intestinal inflammation, typically associated with high levels of intestinal fibrosis [17, 18]. For this aim, we induced chronic colon inflammation and fibrosis in WT and *Ch25h^-/-^* littermate mice with four cycles of 7 days 2.5% DSS in drinking water followed by a 10-day recovery period with normal drinking water. Collagen deposition was determined by Sirius red staining and analysis under transmission light microscopy (Figure 2A). The collagen layer was significantly thinner in *Ch25h^-/-^* mice compared to WT littermate controls (Figure 2B, water animals: WT: 8.5 μm ± 2.1, *Ch25h^-/-^:* 8.7 μm ± 0.8 n.s., DSS animals: WT: 22.4 μm ± 7.2, *Ch25h^-/-^*: 11.7 μm ± 2.6, p= 0.008). Reduced collagen deposition in *Ch25h* knockout mice was confirmed by automated quantification of the collagen layer area (Figure 2C). Additionally, mRNA expression levels for fibrosis markers such as *Tgf-β*, collagen type 3 (*Col-3*) and *Timp-1* were significantly lower in the colon of *Ch25h^-/-^* animals and a clear trend for lower mRNA expression of collagen type 1 *(Col-1*) and lysyl oxidase homolog 2 (*Loxl-2*) in CH25H-deficient mice was found (Figure 2D-H). Expression levels of *Ch25h* were increased in DSS treated animals compared to water controls (Figure 2I).

**Figure 2:**
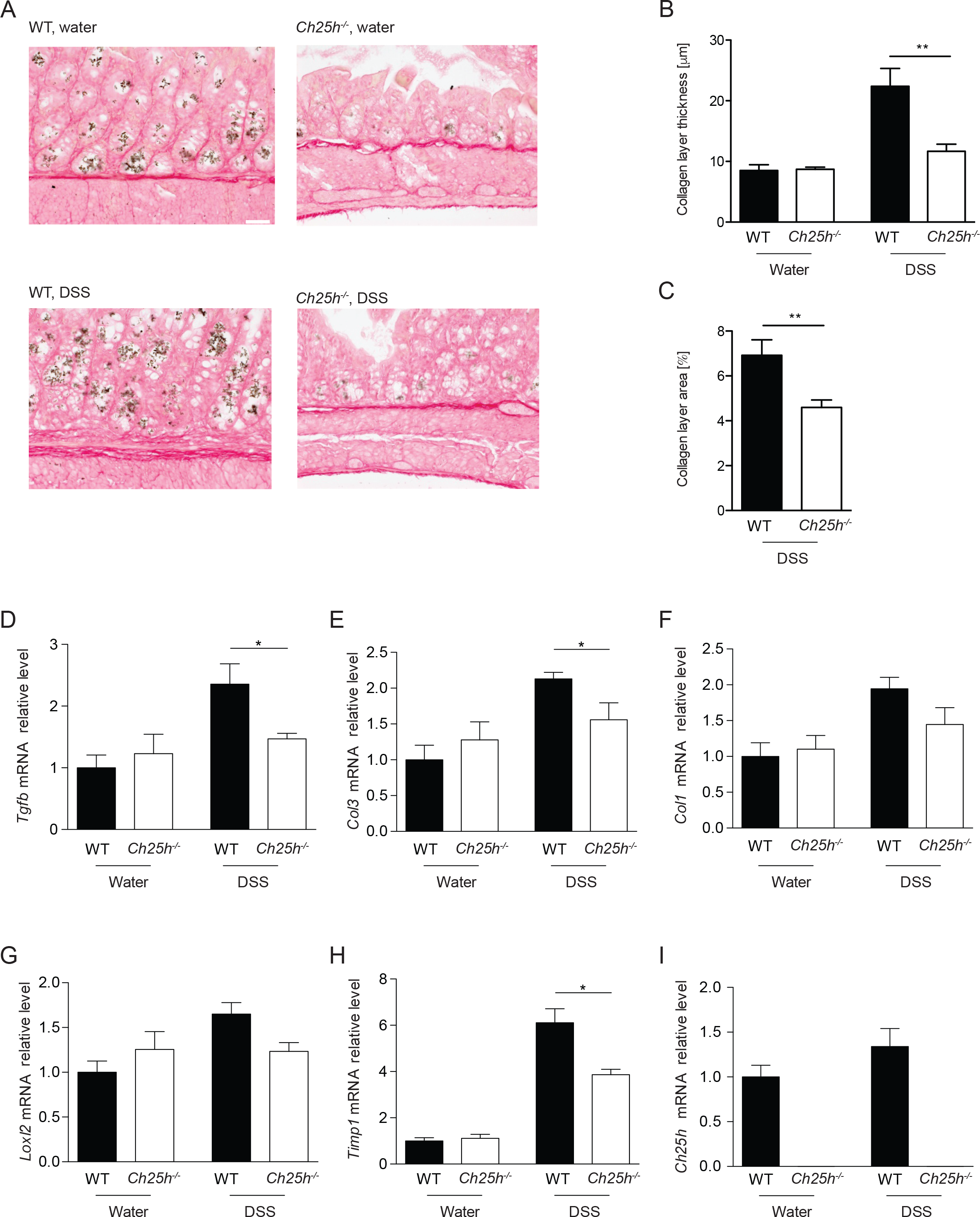
Reduced fibrosis in *Ch25h^-/-^* mice in chronic DSS colitis. *Ch25h^-/-^* and WT female mice were treated for four cycles with 2.5% DSS or water (controls). **(A)** Representative transmission light images of Sirius-red stained intestinal sections of WT and *Ch25h^-/-^* DSS treated mice and water littermate controls. Scale bar: 50 μm. **(B)** Collagen layer thickness calculated from ≥8 positions per graft in representative areas of Sirius-red stained slides with transmission light at 200-fold magnification. **(C)** Quantification of collagen layer area in DSS treated animals using customized Matlab scripts. The colon was analysed for mRNA expression of **(D)** *Tgf-beta*, **(E)** *Col3*, **(F)** *Col1*, **(G)** *Loxl2*, **(H)** *Timp1* and **(I)** *Ch25h* (normalized to *Gapdh*). Expression levels are normalized to water-treated wildtype controls. Statistical analysis: Mann-Whitney U test; * = p<0.05. n = 4-6 per group.

Of note, thinner collagen layer and lower expression levels of fibrosis markers (*Tgf-β*, *Col-3* and *Timp-1)* upon *Ch25h* knockout were not due to reduced inflammation: when colon inflammation was analysed in H/E stained colon sections, the histology score quantifying the inflammatory infiltrate and the epithelial damage, was even higher in *Ch25h^-/-^* mice compared to WT littermates (Figure 3A). Further, macroscopic aspects of intestinal inflammation such as the murine endoscopic index of colitis severity (MEICS) and spleen weight did not differ between both genotypes (Figure 3B). In summary, in chronic DSS colitis, intestinal collagen deposition was reduced in the absence of CH25H, independent from effects of CH25H knockout on intestinal inflammation.

**Figure 3:**
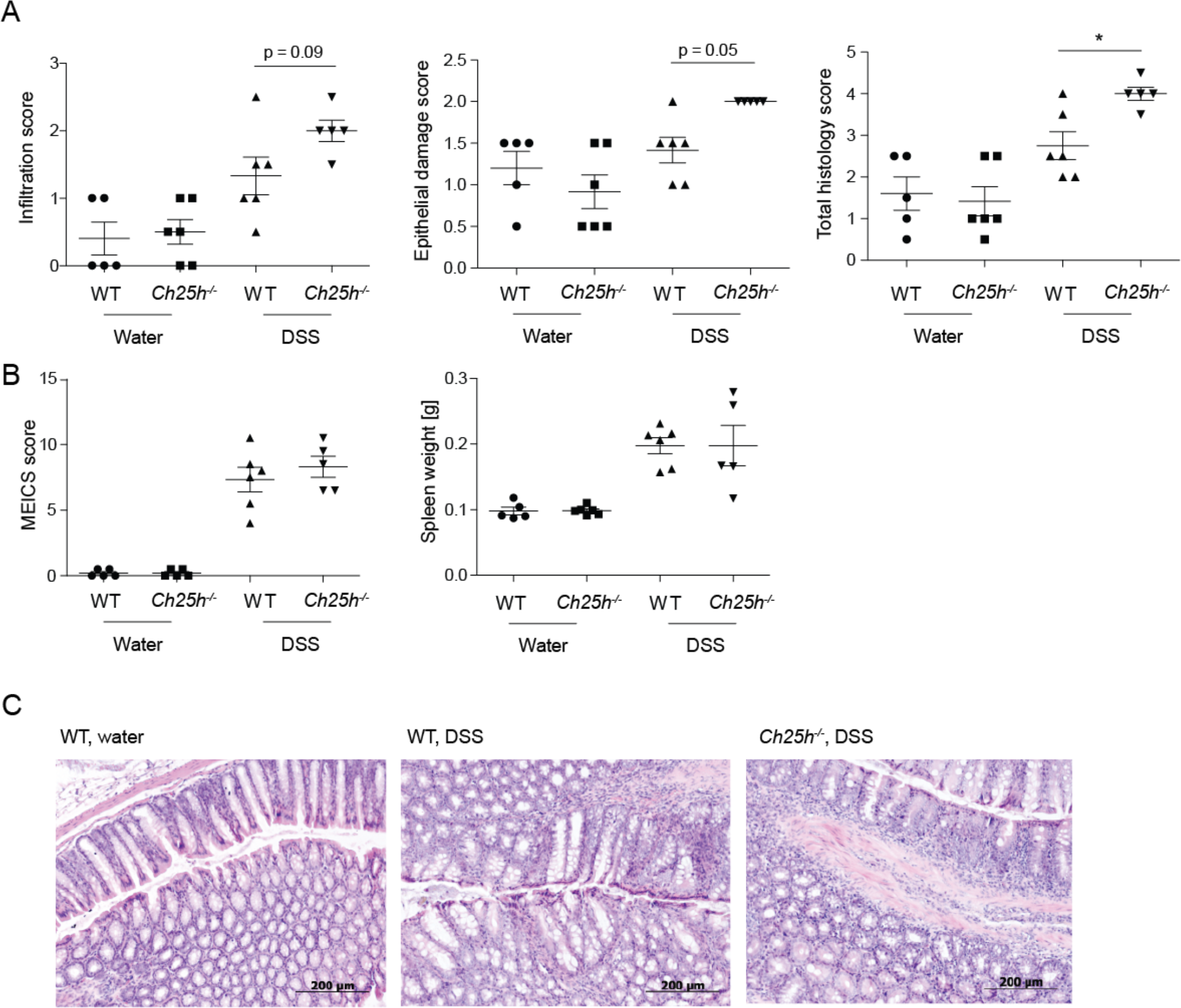
Reduced levels of intestinal fibrosis in CH25H deficient mice is not due to reduced inflammation in chronic DSS colitis. Analysis of colon inflammation in H/E stained colon sections. **(A)** Score of the inflammatory infiltrate (left panel), score for epithelial damage (middle panel) and total histology score (sum of both partial scores, right panel). **(B)** Murine endoscopic index of colitis severity (MEICS) score (left panel) and spleen weight (right panel). **(C)** Representative H/E-stained sections of the distal colon of water control mice (left panel) and DSS treated mice. DSS, dextran sodium sulphate; H/E: hematoxylin and eosin.

### Reduced intestinal fibrosis in the absence of CH25H in a heterotopic transplant model of intestinal fibrosis

To confirm a role of CH25H in intestinal fibrosis in an inflammation-independent model, we employed a recently developed heterotopic transplant model of intestinal fibrosis[19, 20]. Sections of small intestine from either CH25H knockout (*Ch25h^-/-^*) mice or their wildtype littermate controls (WT), were transplanted subcutaneously into the neck of recipient mice of the same genotype[19, 20]. Non-transplanted small bowel sections from *Ch25h^-/-^* mice and WT littermates were used as controls (day 0). Seven days after surgery, the intestinal grafts were collected for analysis (day 7) and collagen deposition was determined by Sirius red staining under transmission light microscopy. At baseline (day 0), cross sections of WT and *Ch25h^-/-^* were histologically indistinguishable with intact epithelial crypts and a thin collagen layer. 7 days post-transplantation, destruction of intestinal epithelial layer, occlusion of the intestinal lumen and a significantly thicker collagen layer was observed[19, 20] (Figure 4A).

**Figure 4:**
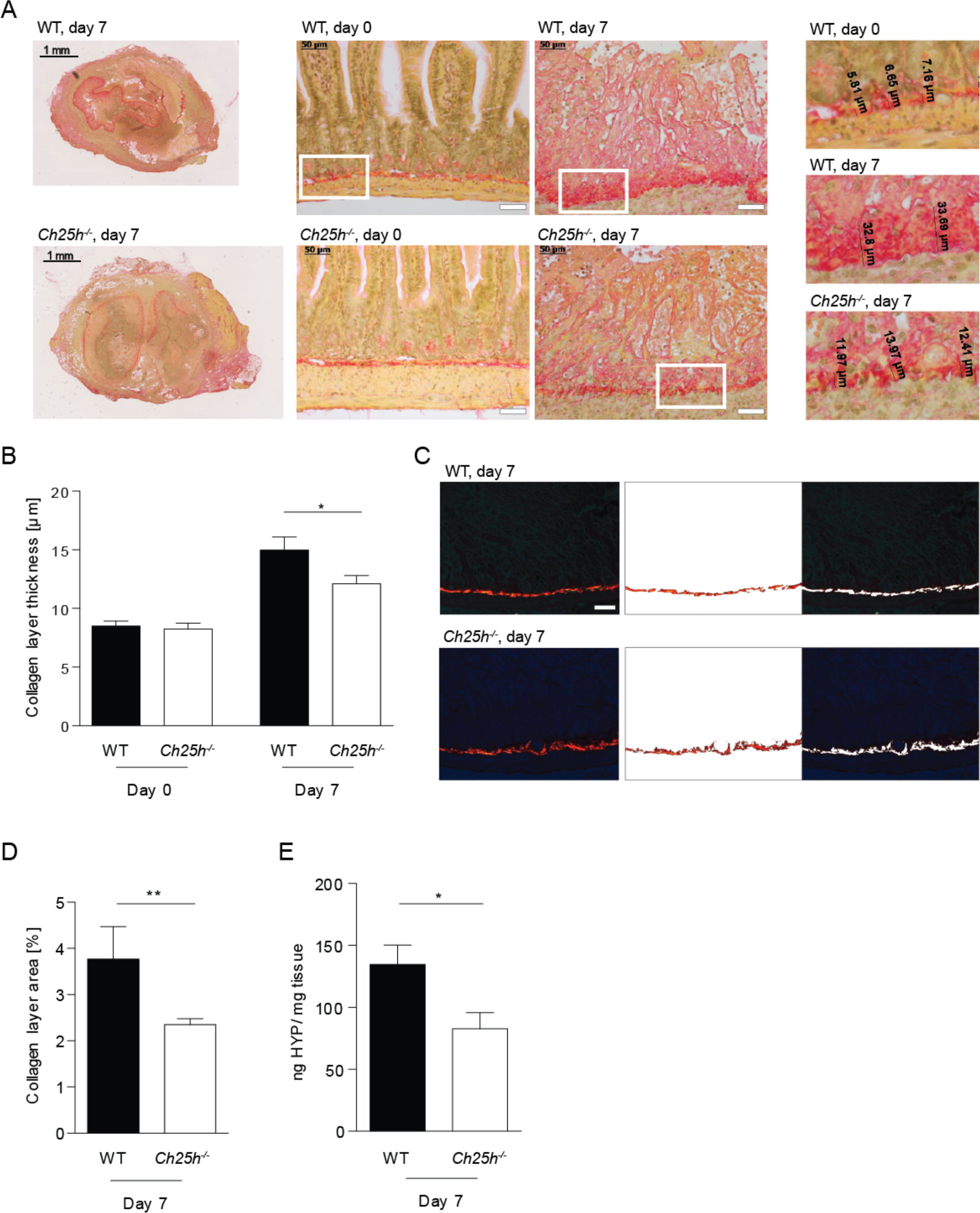
Reduced levels of intestinal fibrosis in CH25H deficient mice in the heterotopic transplantation model. Wildtype and *Ch25h^-/-^* animals were tested in a heterotopic transplantation model for intestinal fibrosis. **(A)** Left panels: Overview (low resolution image) of Sirius red-stained intestinal grafts of WT and *Ch25h^-/-^* mice at day 7 after transplantation. Scale bar: 1 mm. Middle panels: Representative transmission light images demonstrating increased collagen layer thickness in grafts at day 7 compared to freshly isolated intestines at day 0. Upper panels: WT littermate controls. Lower panels: *Ch25h^-/-^*. Scale bar: 50 μm. Right panels: High resolution inserts illustrating measurements of collagen layer thickness. **(B)** Collagen layer thickness calculated from ≥8 positions per graft in representative areas of Sirius-red stained slides with transmission light at 200-fold magnification. **(C)** Image analysis for identification of collagen layer areas using Matlab custom made scripts. Left panel: Original polarized 200x light microscopy image. Middle panel: Collagen layer area. Right panel: Remaining non-collagen tissue. Scale bar: 50 μm. **(D)** Quantification of collagen layer area at day 7 post transplantation using the same strategy as in (C). **(E)** Collagen quantification with hydroxyproline assay. Day 0, freshly isolated intestine. Day 7, intestine 7 days post transplantation. n_WT_ day 0 = 3, n_KO_ day 0 = 9, n_WT_ day 7 = 8, n_WT_ day 7 = 11. Statistical analysis: Mann-Whitney U test; *: p<0.05, **: p<0.01. Bars indicate mean ± SEM. WT, wildtype. CH25H, cholesterol 25 hydroxylase. HYP, hydroxyproline.

The development of intestinal fibrosis was significantly reduced in mice deficient for CH25H indicated by a significantly thinner collagen layer compared to WT littermate controls (Figure 4A, B; day 0: WT: 8.5 μm ± 0.7, *Ch25h^-/-^:* 8.3 μm ± 1.5 n.s., day 7: WT 15.0 μm ± 3.1, *Ch25h^-/-^*: 12.1 μm ± 2.3, p= 0.01). A thinner collagen deposition in *Ch25h* knockout mice was confirmed by polarized light microscopy with automated image analysis and quantification of the collagen layer area (Figure 4C, D). Furthermore, concentration of the collagen metabolite hydroxyproline was significantly lower in *Ch25h* knockout intestinal transplants compared to WT littermate controls (Figure 4E).

*Ch25h* mRNA expression was significantly increased in fibrotic small bowel resections 7 days after transplantation compared to freshly isolated intestine (Figure 5A). Similarly, fibrosis markers including *Col-1* and *Col-3, Timp-1* and *Loxl-2* were induced 7 days after transplantation compared to day 0 (Figure 5B-D). *Ch25h^-/-^* animals displayed a non-significant trend for reduced expression of fibrosis markers compared to WT controls at day 7 (Figure 5B-D). TGF-β protein levels were decreased in *Ch25h* knockout as compared to WT mice, in line with reduced stimulation of profibrotic pathways upon CH25H deficiency (Figure 5E-F). Thus, in agreement with the DSS-induced chronic colitis model, our heterotopic transplant model confirms reduced intestinal fibrosis in the absence of CH25H.

**Figure 5:**
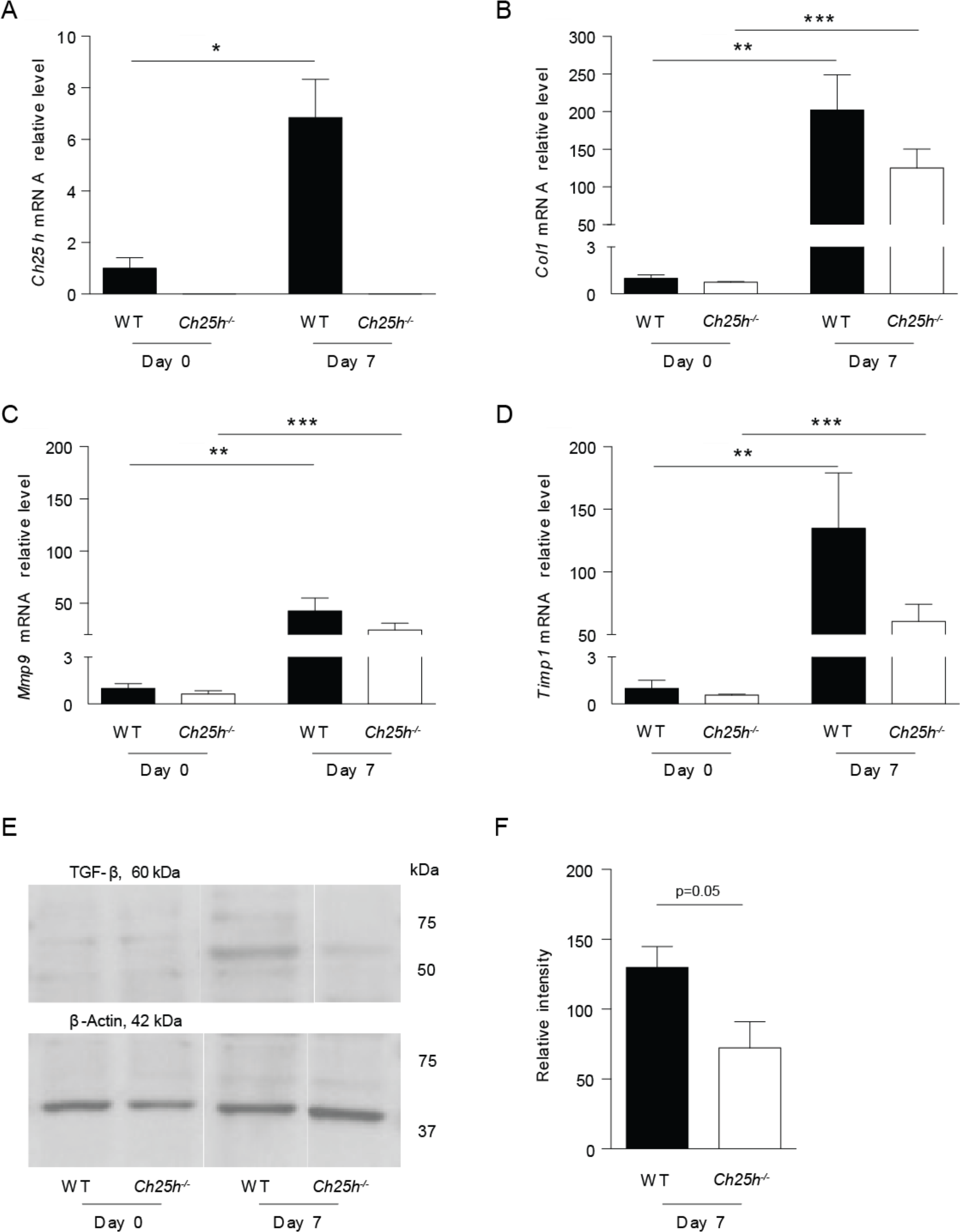
Expression of intestinal fibrosis markers in wildtype and *Ch25h^-/-^* mice. Wildtype and *Ch25h^-/-^* mice were tested in a heterotopic intestinal transplant model. Freshly isolated intestines (day 0) and grafts 7 days after transplantation were analysed for mRNA expression of **(A)** *Ch25h*, **(B)** *Col1*, **(C)** *Mmp9* and **(D)** *Timp1* (normalized to *Gapdh)*. **(E, F)** Analysis of protein expression of TGF-β by Western blot. Expression levels are normalized relative to freshly isolated intestine at day 0 and shown as mean ± SEM. Statistical analysis: A-D: Mann-Whitney U test; * = p<0.05, ** = p<0.01. n_WT_ day 0 = 3, n_KO_ day 0 = 9, n_WT_ day 7 = 8, n_WT_ day 7 = 11. E-F: n = 4, Unpaired t test.

### Recruitment of immune cells into fibrotic small intestine in wildtype and *Ch25h^-/-^*animals

To address changes in immune cells infiltrating the intestinal grafts, lamina propria mononuclear cells were isolated from the grafts 7 days after surgery and an explorative mass cytometry (Cytometry by Time of Flight, CyTOF) analysis with a broad marker panel (Table 2) was performed. Cells were automatically clustered based on similarity of surface marker expression. The immune cell infiltrate was dominated by neutrophils, with a lower fraction of T cells, dendritic cells, monocytes and NK cells (Figure 6A, B). No significant differences between WT and *Ch25h^-/-^* animals were detected for the investigated immune cell populations (Figure 6C). Additionally, histological analysis of IL-17 revealed no differences in IL-17 expression between WT and *Ch25h^-/-^* grafts (Supplementary Figure 1).

**Table 2:**
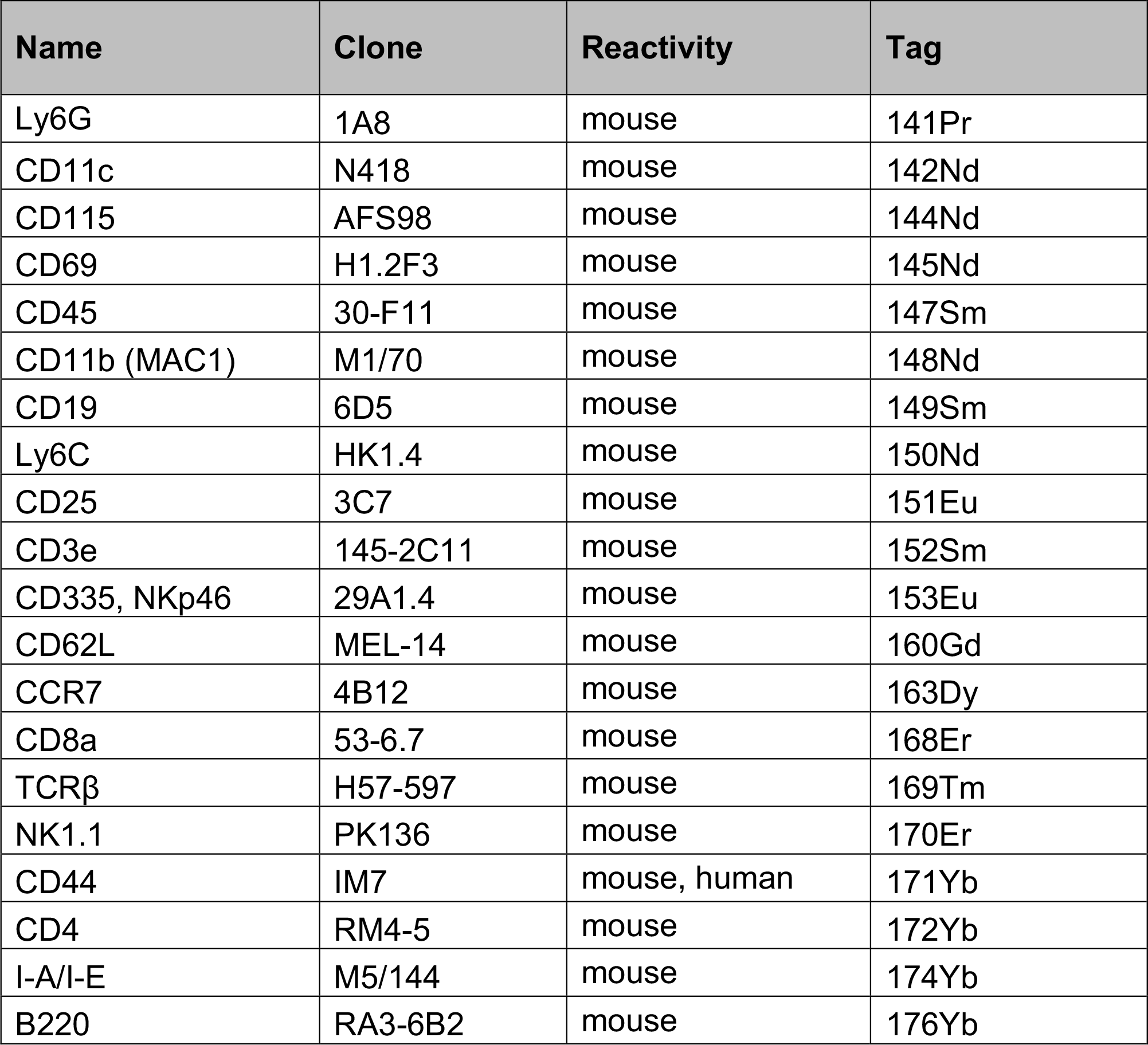
Antibodies used for CyTOF analysis. All antibodies were pre-labelled (Fluidigm).

**Figure 6:**
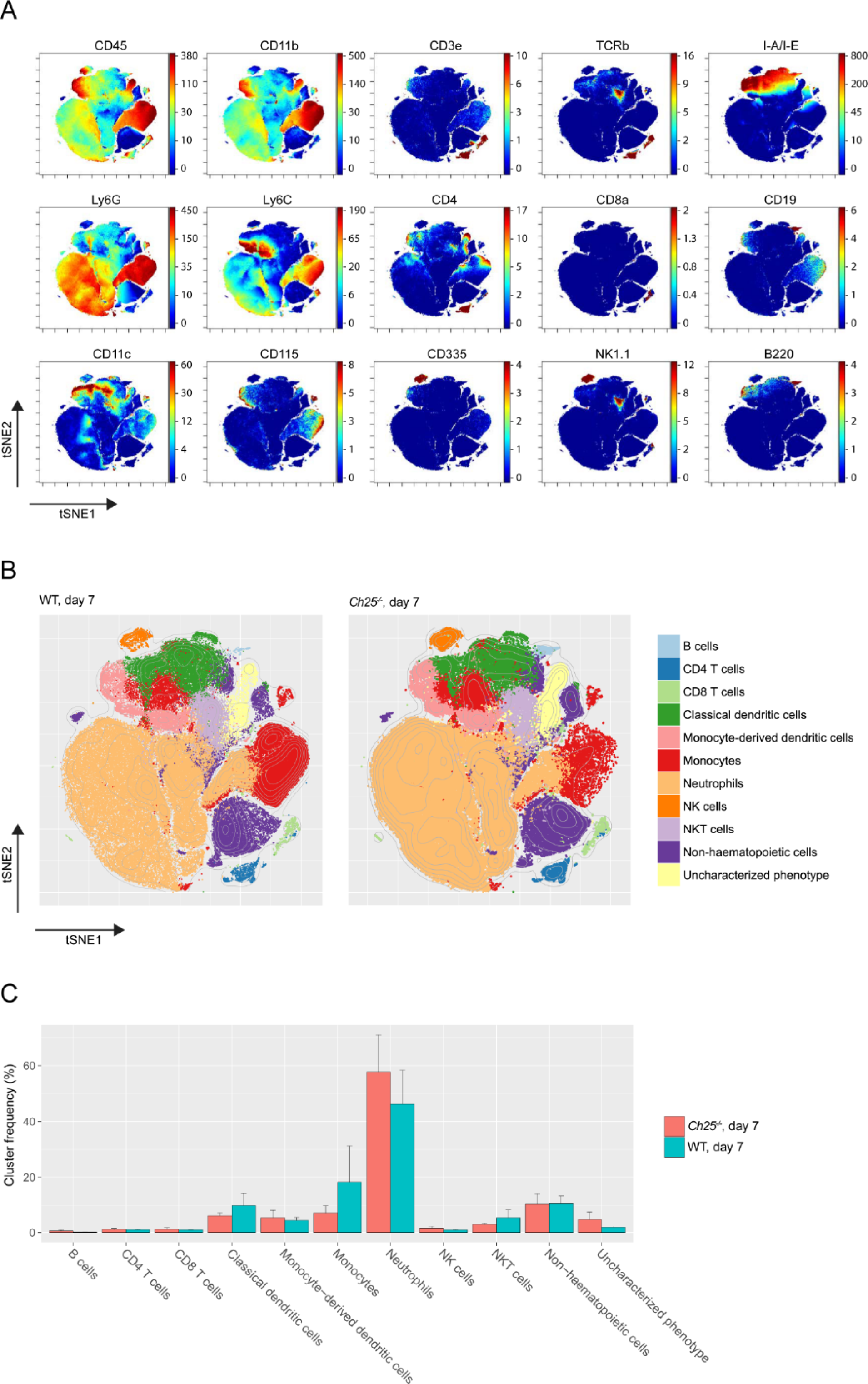
Cells infiltrating the graft do not differ between wildtype and CH25H-deficient mice. Lamina propria infiltrating cells from grafts of wildtype and Ch25h^-/-^ mice were harvested 7 days after surgery and analysed by CyTOF. **(A)** Dimensionality-reduced projection of the entire phenotypical landscape was calculated using the tSNE algorithm with Barnes-Hut approximation (bhSNE). The color-coding represents staining intensity of the specified marker. **(B)** t-SNE maps of each experimental group; 250’000 randomly selected points are plotted. Overlaid in color are cluster designations computed by the Phenograph clustering algorithm. The represented clusters were manually constructed by merging the initial cluster output based on phenotypical similarity until the final number of 11 identifiable clusters was reached. **(C)** Bar plot showing the mean cluster frequencies and error bars representing standard error of the mean (SEM). nWT = 5, nKO = 5

To determine the location of *Ch25h* mRNA expression in the small intestine, RNA *in situ* hybridization using fixed-frozen sections of intestinal grafts and freshly isolated intestines was performed. *Ch25h* mRNA expression was detected in freshly isolated intestines from WT mice and the graft at 7 days after transplantation (Figure 7A, B, Supplementary Figure 2). The *Ch25h* signal appears to be cytoplasmic with the formation of small clusters. *Ch25h* expression was observed in fibroblasts demarcating the necrotic former mucosa layer, but remains of epithelial crypts were not found in the demarcation zone. The *Ch25h-*expresing, spindle shaped fibroblasts are arranged in a band-like fascicle with large ovoid nuclei exhibiting a thinly dispersed chromatin structure and a delicate nuclear membrane without the indentations typically found in the nuclei of histiocytes (Figure 7B, arrows). In contrast, *Ch25h* is not expressed in the inflammatory infiltrate, which mainly consists of neutrophils showing characteristic segmented nuclei (Figure 7B, double arrows). No *Ch25h* expressing lymphocytes were found.

**Figure 7:**
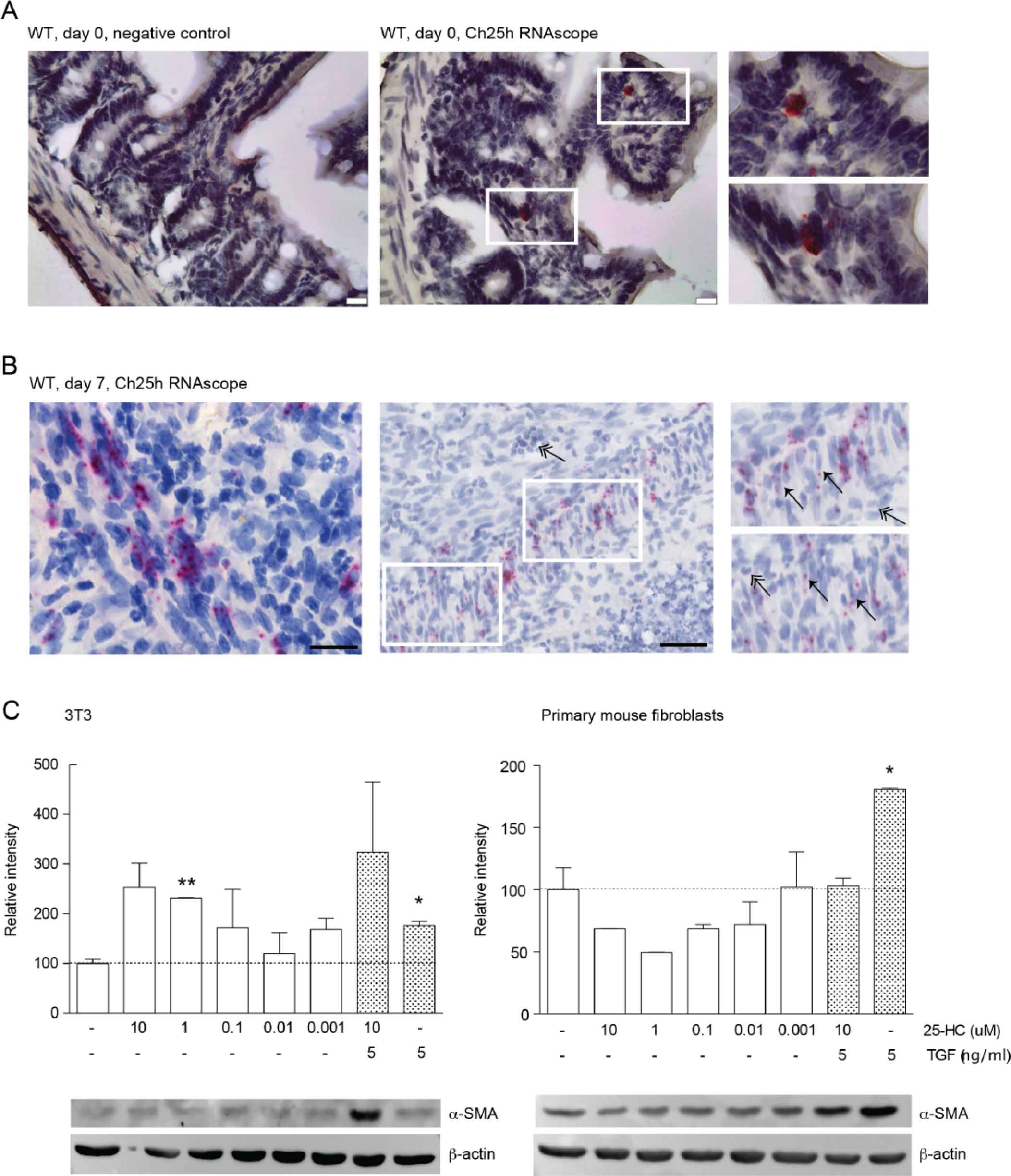
Expression of Ch25h in fibroblasts in intestinal grafts. (A) Representative images of the *in situ* hybridization (RNAscope) analysis of wildtype small intestine. Negative control (a probe for the bacterial gene dihydrodipicolinate reductase, *Dapb*, left panel) and *Ch25h* mRNA (middle panel) are demonstrated with the RNAscope signal shown in red. Right panel: High resolution of inserts of the RNAscope signal. Scale bar: 25 μm. **(B)** Representative images of the *Ch25h* RNAscope analysis of the intestinal grafts of WT mice at day 7 after transplantation, demonstrating accumulation of the CH25H signal in fibroblasts in the former mucosa layer. Scale bar left panel: 20 μm, middle panel: 50 μm. The right panel shows inserts of the middle panel. Fibroblasts are indicated by arrows, neutrophils by a double arrow. **(C)** 3T3 cells (left panel) and primary mouse intestinal fibroblasts (right panel) were treated for 72 h with different concentrations of 25-HC and/or TGF-β as indicated. Samples were analysed for α-SMA protein levels by Western blot. Expression levels are normalized relative to the negative control and shown as mean ± SEM. Statistical analysis: Unpaired t test, the * is relative to the negative control; * = p<.05, **: p<0.01, n = 2.

### 25-HC does not induce myofibroblast differentiation *in vitro*

To address direct effects of the CH25H product 25-HC on fibroblasts, we performed cell culture experiments using primary murine intestinal fibroblasts using a protocol similar to a previous study in a human lung fetal fibroblast cell line (HFL-1) [39]. Addition of TGF-β resulted in increased α-SMA protein expression in intestinal fibroblasts. In contrast, exposure to 25-HC did not affect α-SMA expression at physiological 25-HC concentrations of 0.001-0.1 μM (Figure 7C). Similarly, addition of 25-HC to 3T3 cells did not cause a significant increase of α-SMA expression at concentrations of 0.001-0.1 μM (Figure 7C).

## Discussion

In this study, we demonstrate a role of the oxysterol synthesizing enzyme CH25H in the pathogenesis of intestinal fibrosis. mRNA expression of *CH25H* was upregulated in human intestinal fibrotic tissue of CD patients compared to healthy controls and we found a positive correlation between expression of various fibrosis mediators and *CH25H* expression. Further, we demonstrate a contribution of CH25H to the development of intestinal fibrosis in two murine fibrosis models: in the DSS-induced colitis model, which is commonly used as a model of chronic intestinal inflammation and fibrosis[17, 18], mice lacking the CH25H enzyme showed less collagen deposition and lower mRNA levels of fibrosis mediators. In the recently developed heterotopic transplantation model, lack of CH25H also reduced intestinal collagen deposition as well as levels of the collagen metabolite HYP and the crucial pro-fibrotic factor TGF-β[40, 41].

In several aspects, the newly developed heterotopic transplantation model complements the established DSS-induced chronic model. In the DSS-induced model, fibrosis is induced by repeated disruption of the integrity of the mucosal barrier, resulting in bacterial translocation and lymphocyte infiltration, which promotes chronic colon inflammation. In contrast, in the heterotopic transplantation model, fibrosis is associated with ischemia and hypoxia, independent from inflammatory processes. Here, fibrosis is reliably induced within 7 days after transplantation. This new model reflects important aspects of the human disease such as occlusion of the lumen, expression of TGF-βand α-SMA, as well as collagen deposition in the extracellular matrix[20]. Bacterial translocation and the pathogen associated molecular pattern (PAMP)-associated signalling is not a prerequisite of fibrosis in this model as collagen deposition is also increased following transplantation of small bowel from germfree mice and MyD88 deficient mice (M. Hausmann unpublished observations). In the heterotopic transplant model, fibrosis is also observed in the absence of B and T cells in RAG2 deficient mice (M. Hausmann, unpublished observations). Robust reduction of fibrosis upon CH25H knockout in two independent fibrosis models clearly strengthens the validity of our findings. Our data demonstrate *Ch25h* expression in local fibroblasts in intestinal grafts, which potentially leads to local 25-HC production by fibroblasts. However, it remains unclear, which cell(s) respond to 25-HC during fibrosis induction. In a previous *in vitro* study, 25-HC induced nuclear factor-κB (NF-κB) activation, subsequent release of TGF-β in human fetal lung fibroblasts, ultimately leading to α-SMA expression and myofibroblast differentiation[39]. In support of this finding, activation of NF-κB by 25-HC was also demonstrated in primary rat hepatocytes and in a human monocytic cell line[42, 43]. However, in our study, addition of 25-HC to 3T3 fibroblasts or primary murine intestinal fibroblasts failed to induce α-SMA expression. Therefore, in the intestine, cells different from fibroblasts might be responsible for TGF-β production upon 25-HC exposure.

Previous reports demonstrated that 25-HC inhibits Th17 cell differentiation[24, 44], and *Ch25h* knockout mice have higher numbers of Th17 cells in peripheral lymph nodes and the spleen[45]. Th17 derived IL-17 is a key driver of fibrosis in different organs, including the gut[46-48]. However, histological analysis and quantification of IL-17 revealed no differences between WT and *Ch25h^-/-^* intestinal grafts. In line with these results, the number of CD4 positive cells in grafts from WT and *Ch25h* knockout animals was similar. Overall, the inflammatory infiltrate in intestinal grafts could not be distinguished between mice of both genotypes, which is well in line with involvement of CH25H in profibrotic pathways independent from intestinal inflammation. Our analysis revealed no decrease in intestinal inflammation between wildtype and *Ch25h^-/-^* littermate controls in chronic DSS colitis (Figure 3) and acute DSS colitis (unpublished observations), which further supports that CH25H affects fibrosis in an inflammation-independent manner.

The CH25H product 25-HC has been shown to modulate several immune responses[21, 23, 25, 28, 29, 33]. 25-HC acts as an acute defence molecule and a master regulator of inflammation by increasing antiviral responses[28-30, 49] and decreasing antibacterial defence mechanisms[33, 44]. Our study suggests an additional role of the oxysterol 25-HC as a mediator of intestinal fibrosis. The induction of wound healing, which potentially leads to fibrosis, might thus start very early in the inflammatory cascade by the acute immune-modulatory activity of 25-HC, which is rapidly induced after an inflammatory stimulus[50]. Despite advances in the treatment of CD associated inflammation, a specific intestinal anti-fibrotic therapy remains an unmet clinical need [7, 8]. Our findings clearly indicate that the hydroxylase CH25H is involved in intestinal fibrosis, making CH25H a potential promising novel therapeutic target to prevent intestinal fibrosis, but additional studies are required to elucidate the exact mechanism how CH25H promotes fibrosis.

## Materials and Methods

### Human tissue from patients with CD and controls

Intestinal tissue was obtained from patients with CD undergoing ileocecal resection due to stenosis in the terminal ileum. Non-fibrotic samples originate from the margin of the resections and fibrotic samples from the thickened fibrosis-affected region. Healthy control samples were obtained from patients undergoing right-sided hemicolectomy due to adenocarcinoma (non-cancer affected ileal resection margin). Immediately after resection, samples were fixed in Tissue-Tek® (O.C.T. Compound, Sakura® Finetek), frozen in isopentane on dry ice and stored at −80°C for RNA extraction.

### Animals

CH25H-deficient mice (*Ch25h^-/-^*) in a C57BL/6 background were kindly provided by Novartis Institutes for BioMedical Research[33] and bred in our animal facility with C57BL/6 mice to generate *Ch25h^+/^-* mice. *Ch25h^+/^-* were then crossed to obtain *Ch25h^-/-^* and *Ch25h^+/+^* (wildtype) littermates. The animals received standard laboratory mouse food and water *ad libitum*. They were housed under specific pathogen-free (SPF) conditions in individually ventilated cages. 7- to 10-week old female littermates were used for all studies.

### DSS-induced chronic colitis

DSS-induced chronic colitis was induced by administration of 4 cycles of treatment with DSS (MP Biomedicals). Every cycle consisted of 7 days of 2.5% DSS followed by 10 days of normal drinking water. Mice were killed 4 weeks after the last DSS cycle. Colonoscopy was scored using the murine endoscopic index of colitis severity (MEICS) scoring system [51]. Histological scoring was performed on H&E-stained distal colon sections as described previously [51, 52].

### Heterotopic intestinal transplant model

The heterotopic mouse intestinal transplant model is an adaptation of the transplantation model of intestinal fibrosis in rats, which have both been described in detail previsously[19, 20]. Briefly, donor small bowel was resected and transplanted subcutaneously into the neck of a recipient animal of the same gender and genotype. A single dose of Cefazolin (Kefzol, 1g diluted in 2.5 ml aqua dest.) was applied i.p. as infection prophylaxis. The time interval between graft resection and subsequent implantation was less than 15 minutes. No anesthesia-related recipient death, post-transplantation recipient death or evidence of infection was observed in any of the animals. Intestinal grafts were removed seven days after transplantation. At explantation, each graft was divided into three equal segments. One segment was fixed in 4% formalin and prepared for histopathological assessment. The remaining segments were snap frozen in liquid nitrogen and stored at −80°C for RNA extraction and hydroxyproline (HYP) assay, respectively.

For each sample, 10 mg of snap frozen tissue was homogenized with 100 μl of ultrapure water in M tubes (Miltenyi Biotec) using a gentleMACS tissue homogenizer (Miltenyi Biotec). Graft collagen content was evaluated using the HYP Assay Kit (Sigma-Aldrich) according to the manufacturer’s instructions. HYP concentration is determined by the reaction of 4-(Dimethylamino)benzaldehyde (DMAB) with oxidized HYP, resulting in a colorimetric product (560 nm), proportional to the presence of HYP. All samples and standards were run in duplicate and absorbance at 560 nm was detected on a SpectraMax M2 fluorescence microplate reader using SoftMax Pro version 5 Software (Molecular Devices).

### RNA isolation, cDNA synthesis and real-time-PCR

Total RNA was isolated using the RNeasy Plus Mini Kit (QIAGEN). For mouse samples, lysis buffer from the kit was added to snap frozen resections, and samples were shredded in M tubes (Miltenyi Biotec) in a gentleMACS tissue homogenizer (Miltenyi Biotec). For human samples 10 μm thick tissue tek sections, containing the full thickness of the intestinal wall (confirmed by H/E staining), were cut using a cryostat. Sections were dissolved in TRIzol (Invitrogen, Life Technologies). Total RNA was prepared according to the manufacturer’s instructions. On-column DNase digestion with RDD buffer (QIAGEN) was performed for 15 min at room temperature. RNA concentration was determined by absorbance at 260 and 280 nm. Complementary DNA (cDNA) synthesis was performed using a High-Capacity cDNA Reverse Transcription Kit (Applied Biosystems) following the manufacturer’s instructions. Real-time PCR was performed using the TaqMan Fast Universal Master Mix (Applied Biosystems) on a Fast 7900HT Real-Time PCR System and results analysed with the SDS software (Applied Biosystems). The real-time PCR started with an initial enzyme activation step (5 minutes, 95°C), followed by 45 cycles consisting of a denaturing (95°C, 15 seconds) and an annealing/extending (60°C, 1min) step. For each sample triplicates were measured and glyceraldehyde-3-phosphate dehydrogenase (GAPDH) was used as endogenous control. Results were analysed by the ∆∆CT method. All gene expression assays were obtained from Life Technologies.

### Analysis of microscopy images

Sections were examined using an AxioCam HRc (Zeiss) on a Zeiss Axio Imager.Z2 microscope with AxioVision release 4.8.2 software. Collagen layer thickness was measured on Sirius-red stained slides in at least eight fields in representative areas at 100-fold magnification by an investigator blinded to the experiment. For the automated microscopy analysis Sirius Red-stained slides were analysed by bright-field microscopy with an additional polarizing filter. Under polarized light Sirius Red-stained collagen assumes a palette of colours ranging from green to red based on the fibrotic maturation process. The polarized light images were analysed using MATLAB software, version 8.6 R2015b (MathWorks). Customized scripts identified the collagen layer of each image by clustering pixels of similar colours in two clusters using the k-means clustering algorithm.

### Western blot

Tissue was lysed in M-PER cell lysis buffer (Thermo Fisher Scientific). Protein levels were determined by bicinchoninic acid (BCA) assay according to the manufacturer’s instructions and equal amounts of protein were loaded onto SDS/PAGE gels. Western blots were performed using monoclonal rabbit anti-mouse TGF-β antibodies (3711S, Bioconcept, 1:1000), polyclonal rabbit anti-mouse β-actin antibodies (4970, 13E5, Cell Signaling, 1:2000), polyclonal goat anti-mouse α-SMA antibodies (PA5-18292, Thermo Fisher Scientific, undiluted) and the horseradish peroxidase-conjugated secondary goat anti-rabbit antibody (sc-2004, Santa Cruz, 1:2000). Luminescence of Western blots was quantified densitometrically with ImageJ software.

### Mass cytometry analysis

Data were acquired on a CyTOF-2.1 mass cytometer (Fluidigm) with an acquisition flow rate of 0.03 ml/min. The following signal processing settings were used: default thresholding scheme, lower convolution threshold of 800 intensity units (IU), minimum event duration of 8 pushes, maximum event duration of 100 pushes, noise reduction active. All samples were spiked with EQ four-element calibration beads during acquisition (Fluidigm; cat. no. 201078) and resulting FCS (Flow Cytometry Standard) files were normalized with the built-in normalization algorithm (Helios software version 6.5.358) to account for intra- and intersample intensity measurement variability. The data were analysed and visualized with Cytobank software (Cytobank Inc.) and software packages for R programming language flowCore, flowSOM and ggplot2.

### RNA *in situ* hybridization (RNAscope)

*Ch25h* mRNA localization in the murine small intestine was assessed by RNA *in situ* hybridization. Fresh small intestine sections and intestine grafts were harvested and incubated for 24 hours in 4% paraformaldehyde/PBS (PFA/PBS). The PFA/PBS solution was replaced by 10% sucrose in PBS up to the tissue sink to the bottom of the container. This step was repeated with 20% and 30% sucrose solutions and the tissue was embedded in Optimal Cutting Temperature (OCT). Sections (3-4 μm) were prepared on Superfrost microscope slides (Thermo Fisher Scientific, Braunschweig, Germany). The RNA *in situ* hybridization was performed using the RNAscope 2.5 HD assay, Red (Advanced Cell Diagnostics, Hayward, CA, USA) following the manufacturer’s instructions. In brief, slides were rehydrated in PBS and were subjected to pre-treatment solutions using the recommended incubation time and temperature. Next, slides were incubated for 2h with a Ch25h probe designed and provided by the supplier. The tissue and assay quality were tested with a positive control probe Peptidyl-prolyl cis-trans isomerase B (Ppib, data not shown) and a negative control probe for the bacterial gene Dihydrodipicolinate reductase (Dapb). Four signal amplification steps were carried out at 40°C followed by two additional steps at room temperature with the appropriate solutions. The fifth amplification step was extended from 30 min to one hour in order to enhance the chromogenic signal. Detection of chromogenic signals was achieved by using the Fast-Red reagent for 10 min. Slides were counterstained with hematoxylin I and mounted with VectaMount Mounting Medium HT-5000 (Vector Laboratories, Burlingame, CA, USA).

### *In vitro* experiments

3T3 cells were maintained in high glucose Dulbecco’s Modified Eagle Medium (DMEM, Life Technologies) supplemented with 10% fetal calf serum (FCS) and kept at 37°C in a humidified atmosphere containing 5% CO_2_. Murine primary fibroblasts were isolated and cultured as described previously[53]. The isolated cells were cultured in 25 cm^2^ culture flasks (Costar, Bodenheim, Germany) with DMEM containing 10% FCS, penicillin (100 IE/mL), streptomycin (100 g/mL), ciprofloxacin (8 g/mL), gentamycin (50 g/mL), and amphotericin B (1 g/mL) at 37°C in a humidified atmosphere containing 10% CO_2_. Non-adherent cells were removed. Once fibroblasts reached 90% confluence, FCS free DMEM-medium was added and they were starved for 24 h prior to compound treatment. Cells were stimulated by treatment for 72 h with 5 ng/ml TGF-β (130-095-067, Miltenyi Biotec), 0.001-10 μM 25-HC (H1015, Sigma-Aldrich) or a combination of the two compounds as indicated.

### Statistical analysis

Data are presented as mean ±SEM unless otherwise indicated. Significance was assessed using the Mann-Whitney U test or the unpaired t test with p < 0.05 considered statistically significant (***p < 0.001, **p < 0.01, *p < 0.05).

### Study approval

For patient data, written informed consent was obtained for anonymous use of patient data and resected parts of human intestine according to the code of conduct for responsible use of surgical rest material (Research Code University Medical Center Groningen, http://www.rug.nl/umcg/research/documents/research-code-info-umcg-nl.pdf, see Code goed gebruik voor gecodeerd lichaamsmateriaal). Mouse experiments were approved by the local animal welfare authority (Tierschutzkommision Zürich, Zurich, Switzerland; registration number ZH183/2014).

## Acknowledgments

This research was supported by a grant from the Swiss National Science Foundation to BM (Grant No. 32473B_156525); a grant from the Hartmann-Müller Foundation to BM and an IOIBD grant to GR, MH and BM. The funding institutions had no role in study design, analysis, interpretation of the data and writing of the manuscript. The authors would like to thank Silvia Lang and Kirstin Atrott for technical assistance and Matteo Berchier for the help with the image analysis algorithm.

## Author Contributions

TR, MH, AWS, GR and BM conceived, designed, and supervised the study. TR, MH, MNGA, BW, CM, AW and MRS performed experiments and were involved in data analysis. WTvH and GD were involved in acquisition of human data and samples. PHIS and CAW performed in situ hybridization experiments. VT performed the CyTOF analysis. SL was involved in the histological analysis. TR, MH and BM wrote the paper. MS and GR critically revised the manuscript and added important intellectual content. All authors corrected, and approved the manuscript.

---

**Supplementary Figure 1:**
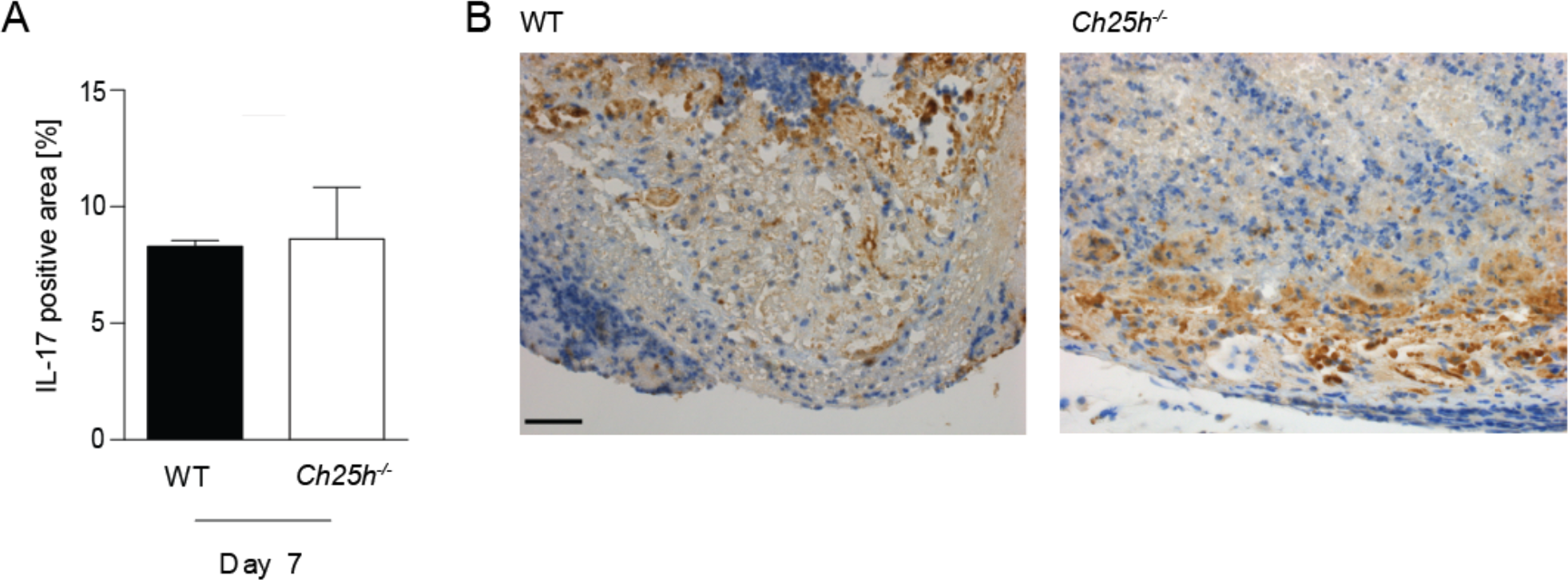
Intestinal grafts from wildtype and CH25H deficient mice do not differ regarding IL-17 expression. **(A)** IL-17 positive areas in intestinal grafts from wildtype and *Ch25h^-/-^* animals identified by automated image analysis using Matlab custom made scripts. The percentage of IL-17 positive staining relative to the graft area is indicated. No significant differences were detected. n_WT_ = 3, n_KO_ = 2, Unpaired t test. **(B)** Representative IL-17 stained pictures; left panel: WT, right panel: *Ch25h^-/-^*. Scale bar: 50 μm.

**Supplementary Figure 2:**
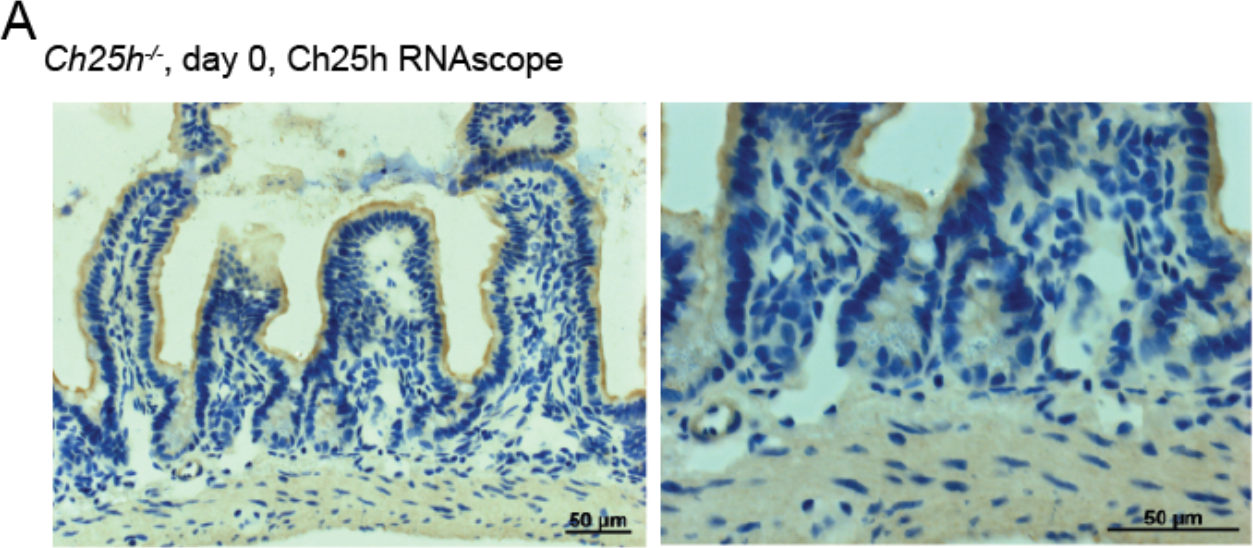
Specificity of *Ch25h* RNAscope staining. **(A)** Freshly isolated intestine of a *Ch25h^-/-^* mouse, demonstrating absence of *Ch25h* mRNA staining.

## Abbreviations

CD: Crohn’s disease
CH25H: cholesterol 25-hydroxylase
COL: collagen
COPD: chronic obstructive pulmonary disease
DSS: dextran sodium sulfate
ECM: extracellular matrix
GAPDH: glyceraldehyde 3-phosphate dehydrogenase
HC: hydroxycholesterol
HYP: hydroxyproline
IBD: Inflammatory bowel disease
IL: interleukin
LPS: lipopolysaccharide
MMP: matrix metalloproteinase
SMA: smooth muscle actin
TGF: transforming growth factor
TIMP: tissue inhibitor of metalloproteinases
TLR: toll like receptor
UC: ulcerative colitis
WT: wildtype

